# Differential Canalograms Detect Outflow Changes from Trabecular Micro-Bypass Stents and Ab Interno Trabeculectomy

**DOI:** 10.1101/044487

**Authors:** Hardik A. Parikh, Ralitsa T. Loewen, Pritha Roy, Joel S. Schuman, Kira L. Lathrop, Nils A. Loewen

## Abstract

The increasing prevalence of glaucoma, a leading cause of blindness, makes the development of safer and more effective treatment more urgent. Recently introduced microincisional glaucoma surgeries that enhance conventional outflow offer a favorable risk profile but can be unpredictable. Two paramount challenges are the lack of an adequate surgical training model for new surgeries and the absence of pre-or intraoperative guidance to sites of reduced flow. To address both, we developed an ex vivo training system and a differential, quantitative canalography method to assess outflow enhancement by trabecular micro-bypass (TMB) implantation or by ab interno trabeculectomy (AIT). TMB resulted in insignificant (p>0.05) outflow increases of 13±5%, 14±8%, 9±3%, and 24±9% in the inferonasal, superonasal, superotemporal, and inferotemporal quadrants. AIT caused a 100±50% (p=0.002), 75±28% (p=0.002), 19±8%, and 40±21% increase in those quadrants. AIT eyes had a 7.5 (p=0.01), 5.7 (p=0.004), 2.3, and 1.8-fold greater outflow enhancement than matching quadrants of paired TMB-implanted eyes. Quantitative canalography demonstrated that TMB, when successful, provided focal outflow enhancements, while AIT achieved a more extensive access to outflow pathways including and beyond the surgical site itself.

## Introduction

Advances in glaucoma surgery device engineering in the sub-millimeter range and improved surgical techniques have caused a rapid evolution of microincisional glaucoma surgeries (MIGS)^1^. Compared to traditional trabeculectomies and tube shunts^2^, these procedures are significantly faster and have a favorable risk profile^1,3^. This also allows to combine procedures^3^ even in complex scenarios^4,5^. The procedures described here, a trabecular meshwork micro-bypass stent (TMB, iStent^®^ G1, Glaukos Corporation, Laguna Hills, CA, USA) and trabectome-mediated ab interno trabeculectomy (AIT, trabectome, Neomedix, Tustin, California, USA) are the only MIGS approved by the Food and Drug Administration of the United States (USFDA). The TMB is a heparin-coated titanium stent measuring 1 x 0.3 mm that is inserted through the trabecular meshwork (TM) into Schlemm’s canal^6^. AIT is a plasma surgery ablation technique that uses a 550 kHz bipolar electrode tip to remove the TM^1^. Ablation is performed over 90 to 180 degrees, allowing to tap into several drainage segments in comparison to single-access MIGS procedures^7^.

The main shortcoming of both procedures is that they are relatively difficult to master and that surgical success of bypassing or removing the TM depends on a functioning downstream collector channel system. There is currently no method to choose the surgical site based on where reduced flow areas are. Similarly, there is no way to directly study where the remaining outflow resistance resides in patients where outflow enhancement fails^3^ despite an otherwise correct surgical technique that should have eliminated the primary resistance of the TM^8,9^. Past studies have quantified the bulk outflow of aqueous humor^10^ or modeled focal outflow mathematically^11^. Tracers, such as cationized ferritin^12^ and fluorescent beads^13,14^, allow to highlight areas of high flow through the TM, but either have cytotoxic effects^15^ or do not easily permit the examination of elements downstream of the TM.

In this study, we developed a quantitative, differential canalography technique to compare conventional outflow enhancement after TMB implantation and AIT. We hypothesized that these procedures would yield distinctly different outflow patterns and developed a novel MIGS training model using enucleated pig eyes.

## Results

Canalograms could be obtained in 41 out of 42 eyes (Fig. 1). Collector channels of the outflow network could be readily visualized using a new two-dye reperfusion technique. The initial filling times for fluorescein (FU) and Texas red (TR) were measured in pilot eyes to determine the proper dye sequence. These times were not significantly different (p=0.06, n=12). A normalization coefficient corrected TR values to match FU at select time points, thereby allowing the comparison of flow rates before and after each intervention. FU demonstrated an average increase of 56±8% fluorescence units compared to TR over 4 quadrants (Supplementary Fig. S1). Eight eyes were used here to achieve >80% power (α=0.05, two-tailed). Significant differences existed in the inferonasal (IN; p=0.028), superonasal (SN; p=0.048), and superotemporal (ST) quadrants (p=0.040). The chromophore fluorescence intensities in the perilimbal region graphed over 15 minutes for each dye showed relatively linear increases in intensity over time as the dyes crossed the TM and the downstream outflow tract (Fig. 2A). The intensity slopes of TR and FU were characteristic for each dye. TR had a lower peak intensity than FU. Dyes did not exhibit chromophore quenching at the concentrations used in our experiments (Fig. 2B).

**Figure 1:**
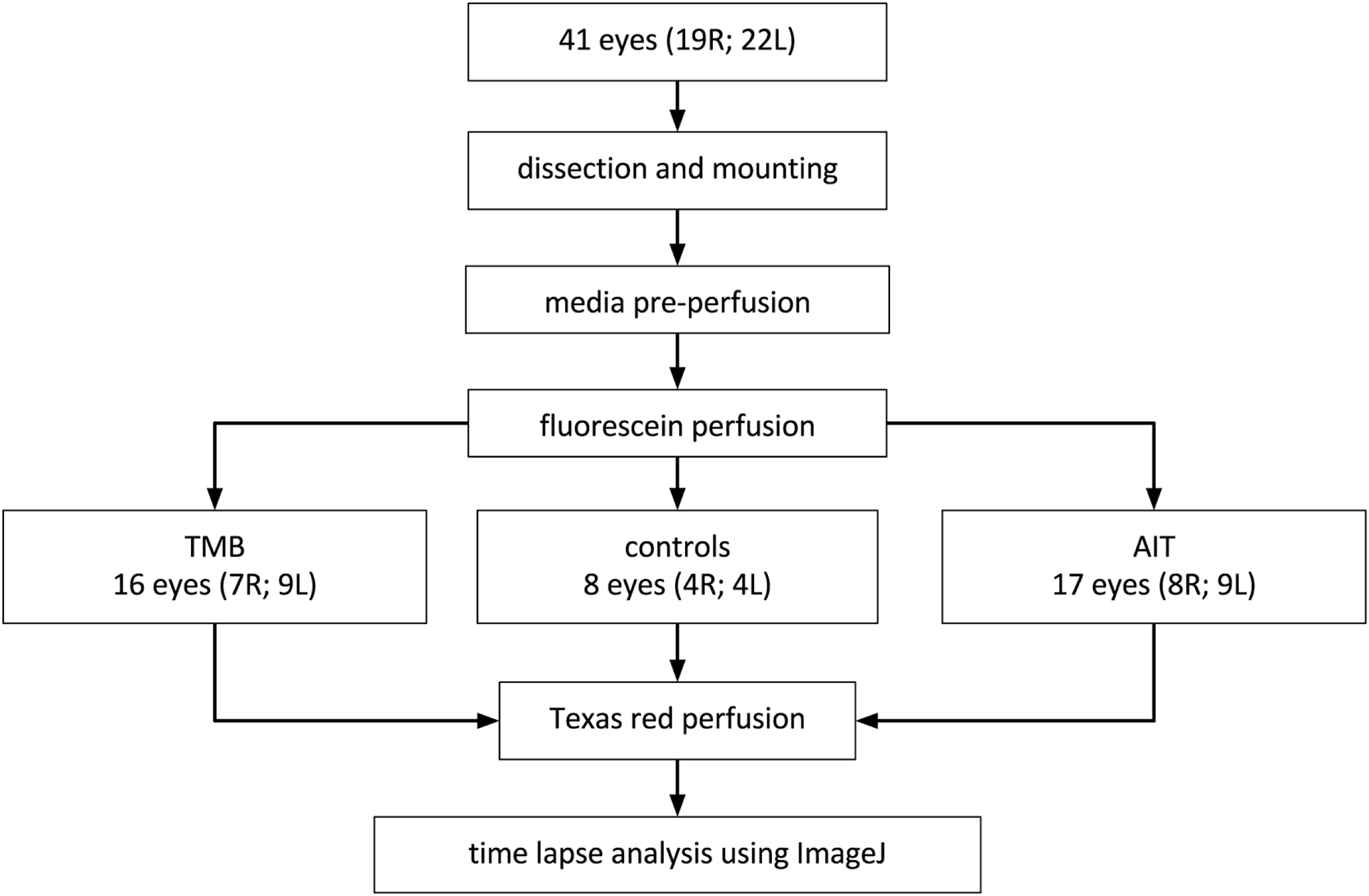
Group assignment. Flowchart detailing assignment to controls, trabecular micro-bypass (TMB) implantation or trabectome-mediated ab interno trabeculectomy (AIT).

**Figure 2:**
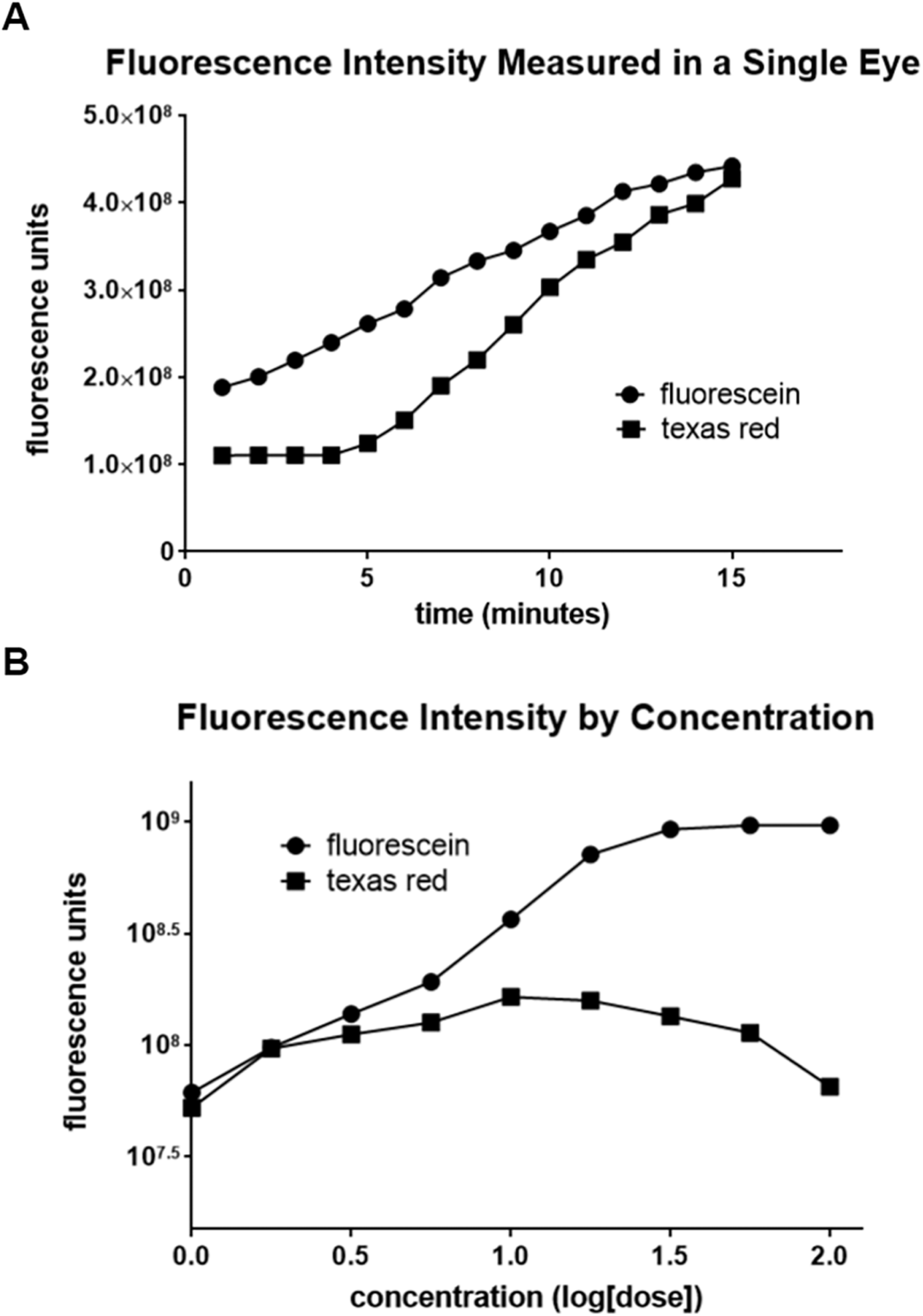
Fluorescein and Texas red fluorescence intensity and concentrations. A) A steeper increase of fluorescence intensity was observed for Texas red than with fluorescein. B) Testing of logarithmic concentrations of fluorescein and Texas red indicated that the concentrations of dyes used for the experiments (concentration=1.0) did not fall under the range of dynamic quenching.

The angle of porcine eyes could be readily visualized by gonioscopy as done in human patients (Fig. 3). TMB implantation proceeded under gonioscopic view using the standard inserter but without viscoelastic (Fig. 3A). AIT could be performed in similar fashion and with tactile feedback that matches human eyes (Fig. 3B).

**Figure 3:**
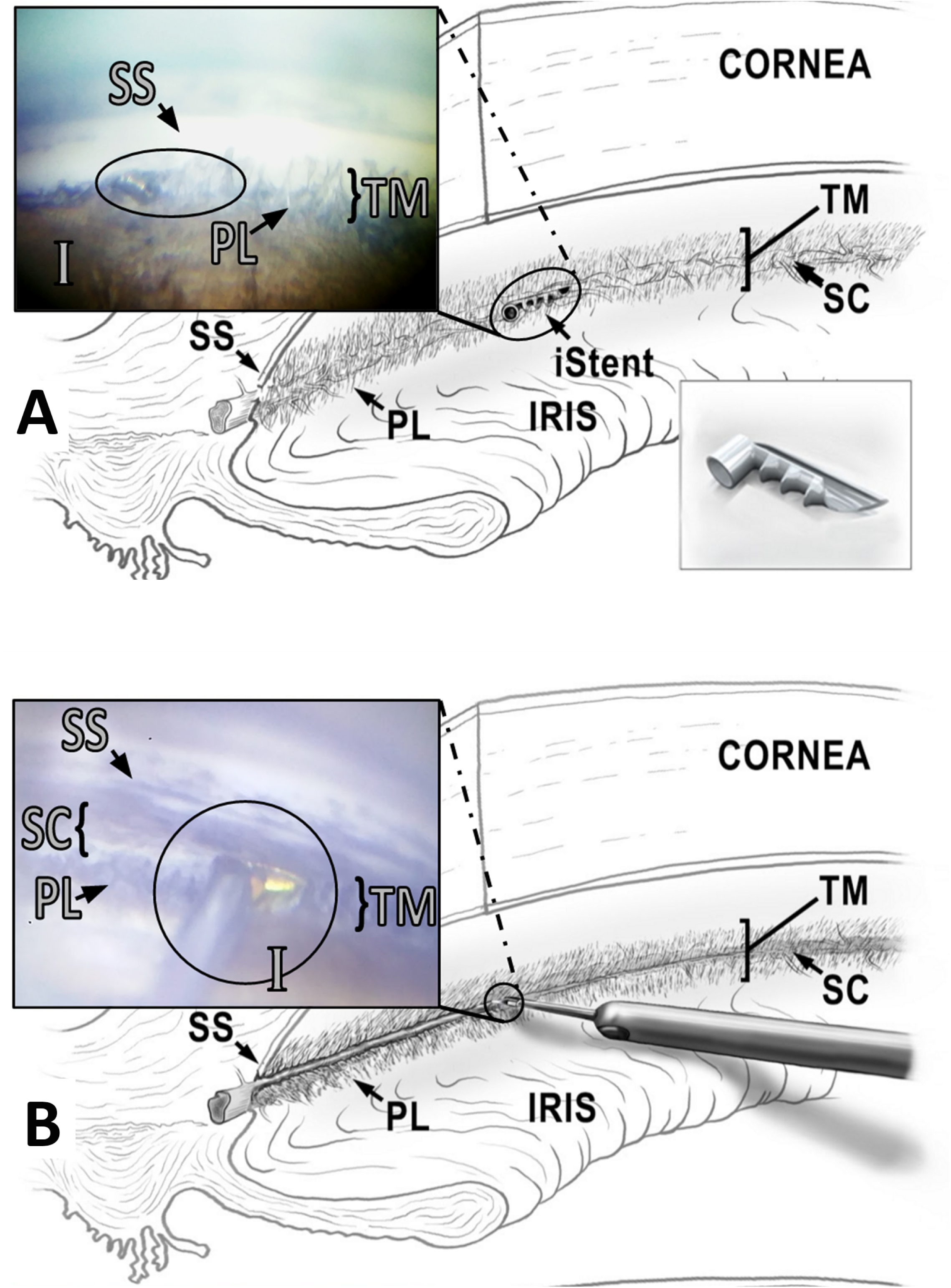
Microincisional glaucoma surgeries in pig eyes. A) Intraoperative view of trabecular micro-bypass (iStent, oval) inserted through the trabecular meshwork and seated within the porcine angular aqueous plexus (distal end) with the snorkel tip parallel to the iris plane (proximal end). B) Intraoperative view of ab interno trabeculectomy (Trabectome, circle). The footplate is visible immediately before trabecular meshwork engagement on the right of the tip. Meshwork has already been ablated on the left of the tip. (SS: scleral spur, TM: trabecular meshwork, I: iris, PL: pectinate ligaments, SC: Schlemm’s canal).

Histology of the angle showed the characteristic pectinate ligaments and the large, wedge-shaped TM of porcine eyes as well as multiple Schlemm’s canal-like structures in variable locations (Fig. 4A). TMB implantation presented as a single lumen that bypassed and displaced the TM and created connections Schlemm’s canal-like structures (Fig. 4B). In contrast, AIT caused a near complete removal of the trabecular meshwork and direct connection to Schlemm’s canal-like structures (Fig. 4C). Thermal, coagulative damage to surrounding structures was mostly absent.

**Figure 4:**
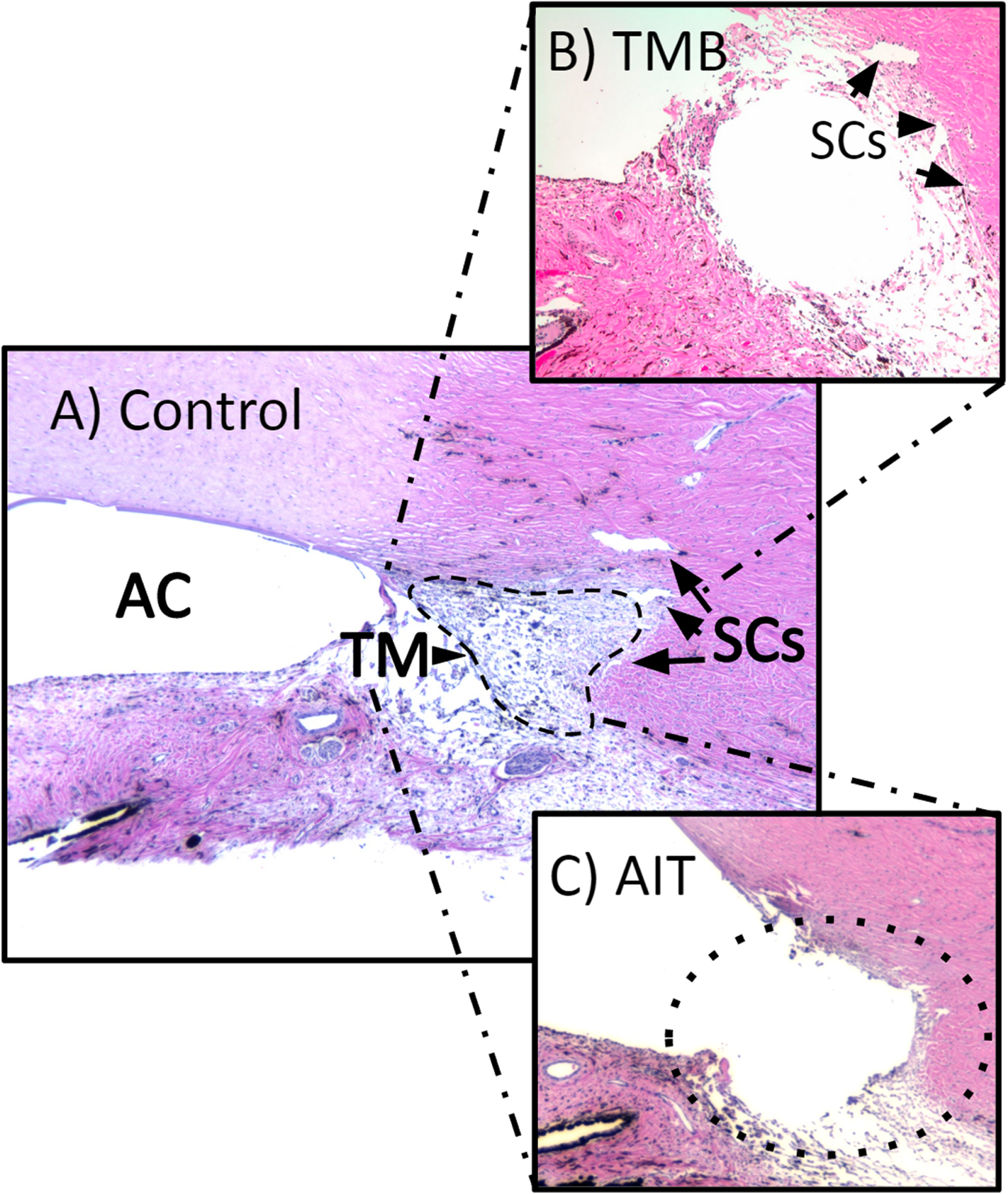
Comparison of sagittal sections of anterior chamber angle. A) Control eye showing anterior chamber with trabecular meshwork (TM) and two Schlemm’s canal-like structures (SC). B) Trabecular micro-bypass (TMB) implantation site is seen as a void. C) Ab interno trabeculectomy (AIT) results in ablation of TM (dotted line).

Time lapses of differential canalograms revealed two different outflow patterns. TMB implantation (Fig. 5A) showed a more focal outflow pattern that extended primarily radially from the location of insertion. The stented quadrant was also usually the first quadrant to show dye filling. In contrast, AIT eyes (Fig. 5D) had increased flow most commonly beginning in the IN or SN, which then extended circumferentially toward centrifugal collector channels. Post-interventional dye filling of collector channels occurred earlier in AIT eyes than in TMB eyes.

**Figure 5:**
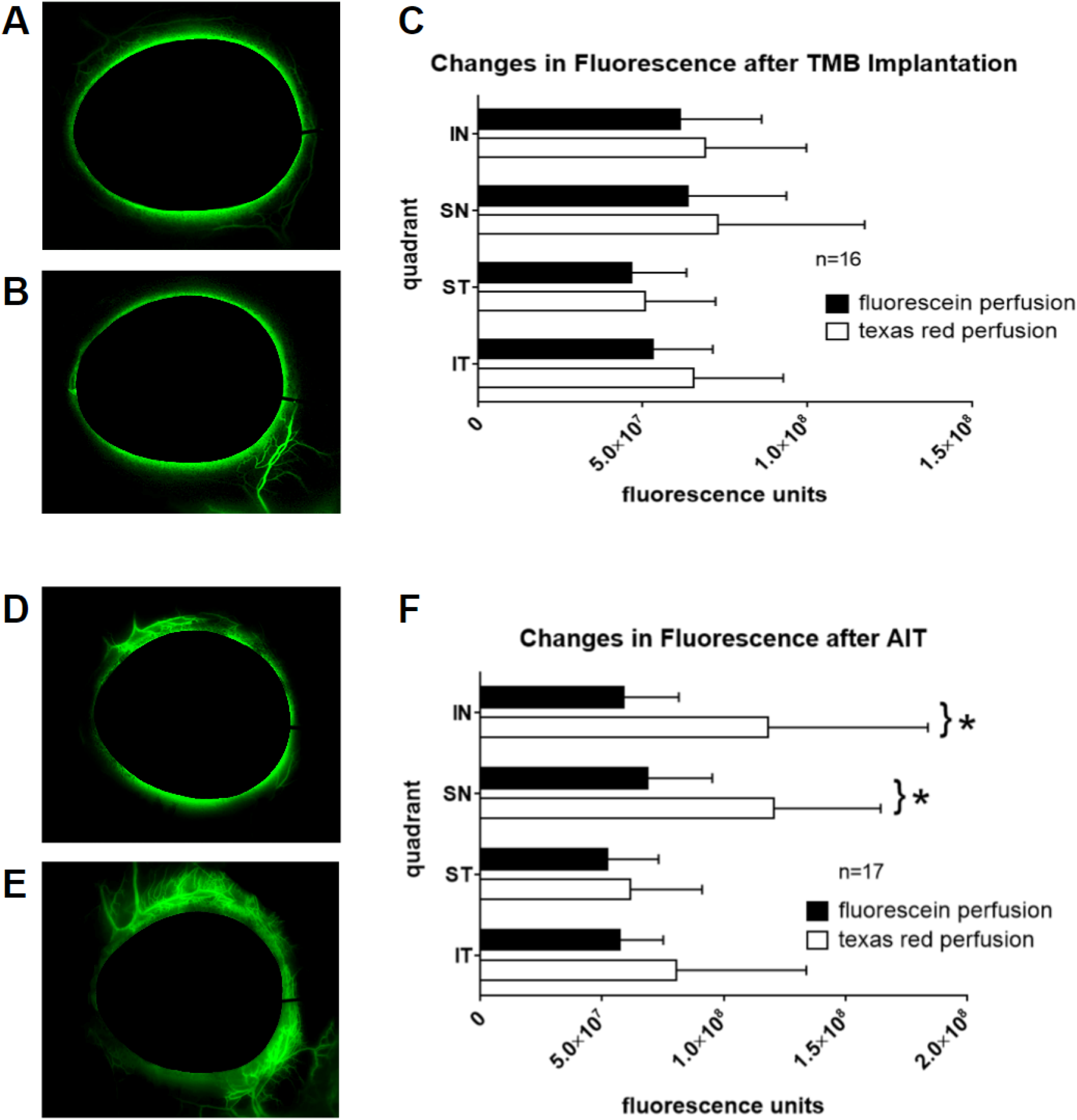
Canalograms and fluorescence intensities after TMB and AIT. Matching canalogram time points before (A) and after TMB (B), which often resulted in a confined, focal outflow enhancement near the implantation site. (C) Average fluorescence increases of 9-24% by quadrant (p>0.05; IN: inferonasal, SN: superonasal, ST: superotemporal, IT: inferotemporal). Canalograms before (D) and after AIT (E), with extensive outflow enhancement beyond ablation ends. (F) Average fluorescence increases of 19-100% by quadrant following nasal AIT (*p<0.05).

Comparison of canalograms before (Fig. 5A) and after TMB implantation (Fig. 5B) in 16 eyes resulted in a statistically insignificant increase of fluorescence averages by 13±5%, 14±8%, 9±3%, and 24±9% in the IN (p=0.25), SN (p=0.44), ST (p=0.51), and IT (p=0.06) quadrants, respectively (Fig. 5C). Outflow facility in these respective quadrants was calculated to be equivalent to 0.82, 0.85, 0.62, and 0.71 microliters per minute before and 0.92, 0.97, 0.68, and 0.88 microliters per minute after that procedure. In contrast, AIT in 17 eyes (Fig. 5D and E) caused fluorescence intensity increases of 100±50% (p=0.0016), 75±28% (p=0.0017), 19±8% (p=0.32), and 40±21% (p=0.13) in the IN, SN, ST, and IT quadrants (Fig. 5F), respectively. Corresponding outflow facility in these quadrants increased from 0.75, 0.87, 0.66, and 0.73 to 1.49, 1.52, 0.78, and 1.02 microliters per minute, respectively. Comparing the two surgical modalities to each other showed that the AIT eyes had a 7.5 (p=0.01), 5.7 (p=0.004), 2.3, and 1.8-fold greater outflow enhancement in the respective quadrants than the TMB eyes (Fig. 6).

**Figure 6:**
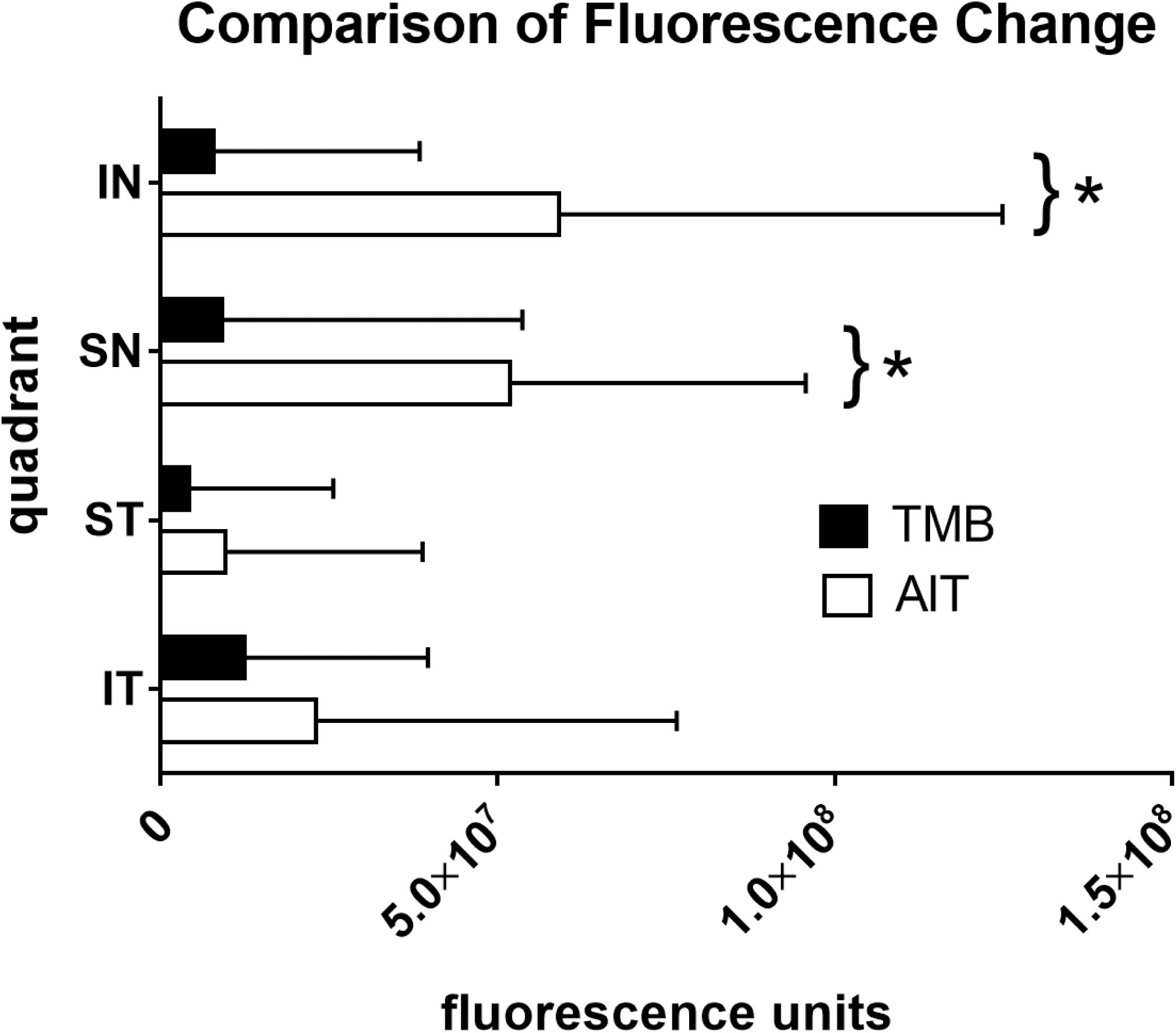
Comparison of postsurgical fluorescence enhancement. Outflow enhancement by ab interno trabeculectomy (AIT) is 1.8 to 7.5 fold larger than that of the trabecular micro-bypass (TMB; *p<0.05; IN: inferonasal, SN: superonasal, ST: superotemporal, IT: inferotemporal).

## Discussion

The trabecular meshwork, a complex sieve-like tissue that permits fluid passage by giant vacuoles, variable pores, and transcytosis^16^, has long been considered to be the principal cause of decreased outflow in primary open angle glaucoma^9^ with most of the resistance thought to be residing in the juxtacanalicular tissue or the inner wall of Schlemm’s canal^17,18^. However, more recent experimental^19^ and clinical^20^ evidence suggests that a large portion of this resistance is located further downstream. Disruption^21^ or ablation of TM^4,20^ would be expected to achieve an intraocular pressure close to that of episcleral venous pressure around 8 mmHg^22^ but this is rarely the case^3^ and failure rates vary considerably from study to study^1,3,20,23^.

Although the procedures discussed here are considered minimally invasive, they are difficult to learn because they are performed on a scale that is approximately 200-fold smaller than that of traditional glaucoma surgery. Maintaining visualization during these procedures is difficult as they require concurrent movement of a surgical goniolens in one hand and MIGS applicators or ablation hand pieces in the other. Because the target tissue, the trabecular meshwork, is in very close proximity to highly vulnerable and well vascularized structures such as the ciliary body band and iris root, novice surgeons can produce serious complications without adequate practice. To address the absence of a microincisional glaucoma surgery model, we created an ex vivo porcine eye system that can also be used to quantify how much outflow improvement was achieved by the trainee using a dye infusion technique. We found that pig eyes provide a highly realistic and inexpensive practice environment that can serve to hone skills before first surgeries in patients. This training model is a powerful preparation tool and does not have to be limited to the procedures discussed here but can also help to master scaffold devices^24^, ab interno sub-Tenon stents^25^, or suprachoroidal shunts^26^, as we can confirm.

Porcine eyes are well suited for this model because they share many features that are similar to human eyes^27^: overall size and shape are comparable; they have a large, wedge-shaped TM that allows TMB and AIT to be performed under the required, direct gonioscopic visualization^1^; possess circumferential drainage segments within the *angular aqueous plexus*^28^ that are considered analogous to Schlemm’s canal^29^; display biochemical glaucoma markers^29^ and giant vacuole formation by Schlemm’s canal endothelium^30^, both of which are seen in human eyes; and present a close match to the human genome^31–33^ which will prove useful in outflow bioengineering approaches^34^.

In order to provide a technique to compare local outflow enhancement from TMB and AIT, we developed a two dye perfusion technique that allows to compare pre-and postsurgical function. The trabecular meshwork and the outer wall of Schlemm’s canal are impermeable to many larger molecules or particles but can be easily passaged by water soluble fluorescent dyes. We selected the organic fluorophores TR and FU because they are readily available and have a very favorable toxicity profile^35^. Spectral domain optical coherence tomography has also recently been used to visualize the aqueous spaces of Schlemm’s canal, the collector channels^36^, and the intrascleral venous network^14^ yet this method does not allow to determine actual flow. Gold nanorods can be used as a Doppler contrast agent to estimate flow^15^ but this method is limited by inflammation^37^.

We had to use a second dye that is different from the first one in postsurgical canalograms because molecules that can flow through the TM may also eventually diffuse into the interstitial space after some time and wash out incompletely. To ensure a valid comparison of pre-and postprocedural canalograms, we thoroughly tested both dyes to account for their characteristic properties. A 19% delay in TR initial filling time in pilot experiments was sufficient evidence for using FU followed by TR for all of the eyes. Choosing this order avoided false positive flow enhancement after the surgical procedures. TR has a molecular weight of 625 g·mol^−1^, almost twice of the molecular weight of FU (332 g·mol^−1^). The two dyes revealed different baseline fluorescence intensities and slope magnitudes, thus resulting in variations in fluorescent intensities at similar time points within both time lapses. Although the relationship between dye concentration and fluorescence intensity is initially linear^38^, very high concentrations of these dyes can result in a decrease of fluorescence intensity as a result of dynamic quenching, an effect described by the Stern-Volmer equation in which excited chromophore molecules will interact with each other and lose energy through processes other than fluorescent emission.^39^ Our testing of logarithmic concentrations of these dyes and measuring their emitted fluorescence through ImageJ ensured that the concentrations used for FU and TR here did not exhibit dynamic quenching. A normalization coefficient corrected TR values to match FU thereby allowing a direct comparison of flow rates before and after each intervention.

The results of this study confirmed our hypothesis that TMB and AIT produce profoundly different outflow patterns. These differences likely correlate with the number of drainage segments that each type of MIGS procedure was able to access effectively. A single point of access to the outflow tract, as delivered by TMB, is thought to enable flow over approximately 60 degrees in human eyes^7^. In contrast, AIT can ablate TM over up to 180 degrees in experienced hands thereby providing access to 180 plus 60 degrees, totaling to 240 degrees of outflow segments^1^. We limited ablation in this study to 90 degrees of the nasal angle because this amount is achievable by most surgeons with ease. It is likely that the porcine eye further highlights the differences between TMB and AIT due to the noncontiguous nature of Schlemm’s canal-like segments in this species. This may limit a successful TMB implantation to less than the 60 degrees of outflow structures in human eyes. Aqueous humor flow rates and facility changes from TMB have been studied in both cadaveric whole eyes^40^ and anterior segments^41^. A single TMB produced a pressure reduction of approximately 6 mmHg. Up until the study presented here, data for trabectome-mediated AIT in animal or cadaveric models has not been available.

In conclusion, we present an ex vivo training model for microincisional glaucoma surgery in pig eyes. We introduce a new differential canalogram technique and find that outflow enhancement of trabectome-mediated ab interno trabeculectomy exceeds that of a trabecular micro-bypass in this species.

## Methods

### Preparation and Pre-Perfusion of the Eyes

Correctly paired, enucleated porcine eyes from a local abattoir were processed within two hours of sacrifice. Each eye was identified as left or right and copiously irrigated with phosphate buffered saline (PBS, Thermo Fisher Scientific, Waltham, MA). Extraocular tissues were trimmed to the length of the globe. Eyes were placed on cryogenic vial cups (CryoElite Cryogenic Vial #W985100, Wheaton Science Products, Millville, NJ) to encompass the optic nerve in a compression-free mount. A 30-gauge needle was inserted through the nasal cornea approximately 2 mm anterior to the limbus parallel to the iris and advanced to the center of the anterior chamber with the bevel up. Eyes were gravity perfused with 37°C clear Dulbecco’s Modified Eagle’s Media (DMEM, Hyclone, GE Healthcare Life Sciences, Piscataway, NJ, USA) for 15 minutes with the pressure set to the regular porcine intraocular pressure of 15 mmHg^27^ as used in anterior chamber perfusion systems^41–43^.

### Chromophore Analysis

When incomplete dye washout was observed in single dye reperfusion pilot experiments, two chromophores with different peak excitation wavelengths (fluorescein (FU): 494 nm, Texas red (TR): 589 nm) were used. The proper detection sensitivities for those were determined using a hemocytometer chamber filled with 10 μL of each dye in clear DMEM at concentrations of 0.0332 mg/mL and 0.28 mg/mL, respectively. Identical average gray values on a 14 bit image were obtained at exposures of 15 and 10 milliseconds, respectively.

Six eyes were first perfused with FU (AK-FLUOR 10%, Fluorescein injection, USP, 100 mg/ml, NDC 17478-253-10, Akorn, Lake Forest, IL) followed by TR (sulforhodamine 101 acid chloride, 10 mg, Alfa Aesar, Ward Hill, MA) to establish whether the order of the dyes would affect the perfusion rate or the intensity of fluorescence; another six eyes underwent the reverse order. Initial filling time, recorded as the time at which the dyes could first be observed entering the proximal outflow tract structures (the perilimbal regions), were recorded for each eye quadrant using ImageJ (ImageJ 1.50b, http://imagej.nih.gov/ij, Wayne Rasband, National Institutes of Health, Bethesda, MD). Eight further control eyes (4 left, 4 right) were run with FU and TR with no intervention in between. After FU and TR time lapse analyses were performed at each eye’s respective half-maximum FU intensity frame, fluorescence units were compared in all four quadrants.

Next, FU and TR were sequentially perfused in a single eye with time lapses (CellSens, Olympus Life Science, Tokyo, Japan) taken as described below. Raw fluorescent intensities of the perilimbal flow patterns were collected every minute for a total of 15 minutes for each dye. When the canalograms of the control eyes revealed consistency between the two chromophores, a normalization coefficient (*c*=1.56) was computed to adjust TR to match FU at a single relative time point in each time lapse pair, allowing for a comparison of flow rates before and after each intervention.

We then determined the fluorescent intensities for FU and TR in each eye at the same relative time points of half-maximum fluorescence of FU as measured in ImageJ. This kept the time factor constant and allowed to compare outflow rates per quadrant in microliters per minute. For this computation, the aqueous humor flow rate of three microliters per minute of porcine eyes was divided by the relative fluorescence of each quadrant measured in control eyes and used as the baseline to compare postsurgical flow.

### Differential Canalograms

The fluorescent tracer reperfusion technique was used in 41 eyes to quantify outflow changes from TMB implantation or AIT. Whole pig eyes were prepared, mounted, and pre-perfused with DMEM as described above. Fluorescein was then gravity-infused at a concentration of 0.0332 mg/ml in clear DMEM for 15 minutes. The chromophore flow pattern was recorded as a time lapse using a stereo dissecting microscope (Olympus SZX16 and DP80 Monochrome/Color Camera; Olympus Corp., Center Valley, PA) equipped for fluorescent imaging (Chroma 49002 GFP cube and Chroma 49004 DSRED cube, Chroma Technology Corp, Bellow Falls, VT). En face images were obtained every 20 seconds at 580 x 610 pixel resolution with 2x2 binning and 14 bit depth (CellSens, Olympus Life Science, Tokyo, Japan). FU infusion was then stopped and the needle removed. A surgeon experienced in microincisional glaucoma surgery (NAL) performed all AITs and TMB implantations as described below (Fig. 3). The incision was sealed in a watertight fashion using cyanoacrylate. Clear DMEM containing TR at a concentration of 0.28 mg/ml was infused for 15 minutes with a time lapse recorded. At conclusion, the eyes were processed for histology. TMB eyes were marked at the site of implantation and the stent was removed. All eyes were rinsed in PBS, hemisected, and fixed with 4% paraformaldehyde at room temperature followed by PBS for 48 hours before being placed in 70% ethanol. A corneoscleral wedge from the surgical site was paraffin-embedded for histological processing, cut at 10 μm thickness, and stained with hematoxylin and eosin (H&E).

### Trabecular Meshwork Bypass Implantation Technique

Sixteen pig eyes underwent TMB implantation (iStent, Glaukos Corporation, Laguna Hills, CA, USA). The surgical technique was analogous to that used in human eyes^44^. With the temporal side of the eye facing the surgeon, a clear corneal incision was created 2 mm anterior to the temporal limbus with a 1.8 mm keratome. The loaded TMB applicator was inserted into the anterior chamber and advanced toward the TM. The stent was driven through the TM using a gentle sweeping motion. Only the proximal end of the stent remained visible in the anterior chamber (Fig. 3A). The stent was ejected from the applicator and the applicator tip was removed. A drop of cyanoacrylate was used to seal the incision.

### Ab Interno Trabeculectomy Technique

After infusion with FU, 17 eyes underwent AIT, performed analogous to AIT in human eyes^1^. Eyes were positioned under a surgical microscope with the temporal side directed toward the surgeon. A 1.8 mm keratome was used to create a clear corneal incision 2 mm anterior to the temporal limbus. The inner third was slightly flared to improve mobility and eliminate striae from torque. The eyes were then tilted by 30 degrees toward the nasal side and a goniolens (Goniolens ONT-L, #600010, NeoMedix Inc., Tustin, CA) was placed on the cornea to visualize the nasal chamber angle. The tip of the trabectome handpiece was inserted into the anterior chamber with constant irrigation, and gentle goniosynechiolysis with the smooth base plate was performed to disinsert pectinate ligaments (Fig. 3B). The TM was engaged and Schlemm’s canal entered with a left and upward movement. TM ablation at 1.1 mW ensued toward the left for 45 degrees with appropriate rotation of the goniolens to maintain visualization. The tip was then disengaged from the TM, rotated 180 degrees within the eye, and positioned at the original starting location. A 45 degree ablation was performed towards the right. The handpiece was removed, and the incision closed watertight with a drop of cyanoacrylate.

### Time Lapse Analysis

Individual time lapses with FU and TR were analyzed using ImageJ software^45,46^. For each fluorescein time lapse, the half-maximum perilimbal fluorescence was calculated, and the appropriate frame containing perilimbal fluorescence that best matched that value was selected for analysis. The same frame number was used in the corresponding Texas Red time lapse for each eye. A masked observer measured raw fluorescence intensities from these frames by outlining the quadrants containing fluorescent outflow channels. Each pig eye was divided into four quadrants: inferonasal (IN), superonasal (SN), superotemporal (ST), and inferotemporal (IT). Quadrant outlines began at the limbus of each quadrant to exclude quantification of fluorescence of the dye in the anterior chamber.

### Statistics

Student’s paired two sample t-test was used to compare outflow changes in the same eyes before and after each intervention. The unpaired t-test was utilized to detect any significant differences between right and left eyes, and postprocedural outflow enhancement between the experimental groups. All t-tests were two-tailed and all data followed a normal distribution. Results were reported as means with standard deviation (mean±SD).

## Acknowledgements

The authors would like to acknowledge Katherine Davoli (Core Grant for Vision Research EY08098) for providing histology. This work was supported by grants from the National Institutes of Health (K08EY022737), Eye and Ear Foundation of Pittsburgh, American Glaucoma Society, Research to Prevent Blindness, and the Alpha Omega Alpha Carolyn L Kuckein Student Research Fellowship.

## Author Contributions

NAL conceived and supervised the study and provided the materials. HAP, RTL, and PR conducted the experiments. HAP analyzed the results. HAP and NAL wrote the main manuscript text. KLL assisted with experimental design and prepared Figure 2. JSS advised on experimental design and critically reviewed the manuscript. RTL greatly contributed to Figure 4. All authors reviewed the manuscript.

## Competing Financial Interests

NAL has received honoraria as a Trabectome trainer for NeoMedix Corporation. NeoMedix Corporation, as well as any of the sponsors aforementioned, have had no role in the design or conduct of this research.

## References

1. Kaplowitz, K., Schuman, J. S. & Loewen, N. A. Techniques and outcomes of minimally invasive trabecular ablation and bypass surgery. Br. J. Ophthalmol. 98, 579–585 (2014).

2. Gedde, S. J. et al. in Am J Ophthalmol 153, 804–814.e1 (2012 Elsevier Inc, 2012).

3. Kaplowitz, K., Bussel, I. I., Honkanen, R., Schuman, J. S. & Loewen, N. A. Review and meta-analysis of ab-interno trabeculectomy outcomes. Br. J. Ophthalmol. (2016). doi:10.1136/bjophthalmol-2015-307131

4. Bussel, I. I., Kaplowitz, K., Schuman, J. S., Loewen, N. A. & Trabectome Study Group. Outcomes of ab interno trabeculectomy with the trabectome by degree of angle opening. Br. J. Ophthalmol. 99, 914–919 (2015).

5. Bussel, I. I. et al. Outcomes of ab interno trabeculectomy with the trabectome after failed trabeculectomy. Br. J. Ophthalmol. 99, 258–262 (2014).

6. Samuelson, T. W. et al. Randomized evaluation of the trabecular micro-bypass stent with phacoemulsification in patients with glaucoma and cataract. Ophthalmology 118, 459–467 (2011).

7. Rosenquist, R., Epstein, D., Melamed, S., Johnson, M. & Grant, W. M. Outflow resistance of enucleated human eyes at two different perfusion pressures and different extents of trabeculotomy. Curr. Eye Res. 8, 1233–1240 (1989).

8. Bill, A. & Svedbergh, B. Scanning electron microscopic studies of the trabecular meshwork and the canal of Schlemm—an attempt to localize the main resistance to outflow of aqueous humor in man. Acta Ophthalmol. 50, 295–320 (1972).

9. Mäepea, O. & Bill, A. Pressures in the juxtacanalicular tissue and Schlemm’s canal in monkeys. Exp. Eye Res. 54, 879–883 (1992).

10. Nau, C. B., Malihi, M., McLaren, J. W., Hodge, D. O. & Sit, A. J. Circadian variation of aqueous humor dynamics in older healthy adults. Invest. Ophthalmol. Vis. Sci. 54, 7623–7629 (2013).

11. Yuan, F. et al. Mathematical Modeling of Outflow Facility Increase With Trabecular Meshwork Bypass and Schlemm Canal Dilation. J. Glaucoma (2015). doi:10.1097/IJG.0000000000000248

12. de Kater, A. W., Melamed, S. & Epstein, D. L. Patterns of aqueous humor outflow in glaucomatous and nonglaucomatous human eyes. A tracer study using cationized ferritin. Arch. Ophthalmol. 107, 572–576 (1989).

13. Hann, C. R. & Fautsch, M. P. Preferential fluid flow in the human trabecular meshwork near collector channels. Invest. Ophthalmol. Vis. Sci. 50, 1692–1697 (2009).

14. Kagemann, L. et al. 3D visualization of aqueous humor outflow structures in-situ in humans. Exp. Eye Res. 93, 308–315 (2011).

15. Wang, B. et al. Gold nanorods as a contrast agent for Doppler optical coherence tomography. PLoS One 9, e90690 (2014).

16. Tamm, E. R. The trabecular meshwork outflow pathways: structural and functional aspects. Exp. Eye Res. 88, 648–655 (2009).

17. Overby, D. R., Stamer, W. D. & Johnson, M. The changing paradigm of outflow resistance generation: towards synergistic models of the JCT and inner wall endothelium. Exp. Eye Res. 88, 656–670 (2009).

18. Grant, W. M. Experimental aqueous perfusion in enucleated human eyes. Arch. Ophthalmol. 69, 783–801 (1963).

19. Schuman, J. S., Chang, W., Wang, N., de Kater, A. W. & Allingham, R. R. Excimer laser effects on outflow facility and outflow pathway morphology. Invest. Ophthalmol. Vis. Sci. 40, 1676–1680 (1999).

20. Parikh, H. A., Bussel, I. I., Schuman, J. S., Brown, E. N. & Loewen, N. A. Coarsened Exact Matching of Phaco-Trabectome to Trabectome in Phakic Patients: Lack of Additional Pressure Reduction from Phacoemulsification. PLoS One 11, e0149384 (2016).

21. Grover, D. S. et al. Gonioscopy-Assisted Transluminal Trabeculotomy, Ab Interno Trabeculotomy. Ophthalmology 121, 855–861 (2014).

22. Sultan, M. & Blondeau, P. Episcleral venous pressure in younger and older subjects in the sitting and supine positions. J. Glaucoma 12, 370–373 (2003).

23. Jea, S. Y., Francis, B. A., Vakili, G., Filippopoulos, T. & Rhee, D. J. Ab interno trabeculectomy versus trabeculectomy for open-angle glaucoma. Ophthalmology 119, 36–42 (2012).

24. Johnstone, M. A. et al. Effects of a Schlemm canal scaffold on collector channel ostia in human anterior segments. Exp. Eye Res. 119, 70–76 (2014).

25. Lewis, R. A. Ab interno approach to the subconjunctival space using a collagen glaucoma stent. J. Cataract Refract. Surg. 40, 1301–1306 (2014).

26. Hoeh, H. et al. Early postoperative safety and surgical outcomes after implantation of a suprachoroidal micro-stent for the treatment of open-angle glaucoma concomitant with cataract surgery. J. Cataract Refract. Surg. 39, 431–437 (2013).

27. McMenamin, P. G. & Steptoe, R. J. Normal anatomy of the aqueous humour outflow system in the domestic pig eye. J. Anat. 178, 65–77 (1991).

28. Tripathi, R. C. Ultrastructure of the exit pathway of the aqueous in lower mammals:(A preliminary report on the ‘angular aqueous plexus’). Exp. Eye Res. 12, 311–314 (1971).

29. Suárez, T. & Vecino, E. Expression of endothelial leukocyte adhesion molecule 1 in the aqueous outflow pathway of porcine eyes with induced glaucoma. Mol. Vis. 12, 1467–1472 (2006).

30. McMenamin, P. G. & Steptoe, R. J. Normal anatomy of the aqueous humour outflow system in the domestic pig eye. J. Anat. 178, 65–77 (1991).

31. Groenen, M. A. M. et al. Analyses of pig genomes provide insight into porcine demography and evolution. Nature 491, 393–398 (2012).

32. Flicek, P. et al. Ensembl 2014. Nucleic Acids Res. 42, D749–55 (2014).

33. Pairwise Alignment Human vs Pig LastZ Results. Ensembl at <http://useast.ensembl.org/info/genome/compara/mlss.html?mlss=716>

34. Loewen, R. et al. A Porcine Anterior Segment Perfusion and Transduction Model with Direct Visualization of the Trabecular Meshwork. Invest Ophthalmol Vis Sci. In press (2016).

35. Alford, R. et al. Toxicity of organic fluorophores used in molecular imaging: literature review. Mol. Imaging 8, 341–354 (2009).

36. Kagemann, L. et al. Visualization of the conventional outflow pathway in the living human eye. Ophthalmology 119, 1563–1568 (2012).

37. Gabriele Sandrian, M. et al. Inflammatory response to intravitreal injection of gold nanorods. Br. J. Ophthalmol. 96, 1522–1529 (2012).

38. Guilbault, G. G. Practical fluorescence. 3, (CRC Press, 1990).

39. Kadadevarmath, J. S., Malimath, G. H., Melavanki, R. M. & Patil, N. R. Static and dynamic model fluorescence quenching of laser dye by carbon tetrachloride in binary mixtures. Spectrochim. Acta A Mol. Biomol. Spectrosc. 117, 630–634 (2014).

40. Hunter, K. S., Fjield, T., Heitzmann, H., Shandas, R. & Kahook, M. Y. Characterization of micro-invasive trabecular bypass stents by ex vivo perfusion and computational flow modeling. Clin. Ophthalmol. 8, 499–506 (2014).

41. Bahler, C. K., Smedley, G. T., Zhou, J. & Johnson, D. H. Trabecular bypass stents decrease intraocular pressure in cultured human anterior segments. Am. J. Ophthalmol. 138, 988–994 (2004).

42. Chowdhury, U. R. et al. ATP-sensitive potassium (KATP) channel activation decreases intraocular pressure in the anterior chamber of the eye. Invest. Ophthalmol. Vis. Sci. 52, 6435–6442 (2011).

43. Bahler, C. K., Fautsch, M. P., Hann, C. R. & Johnson, D. H. Factors influencing intraocular pressure in cultured human anterior segments. Invest. Ophthalmol. Vis. Sci. 45, 3137–3143 (2004).

44. Nichamin, L. D. Glaukos iStent Trabecular Micro-Bypass. Middle East Afr. J. Ophthalmol. 16, 138–140 (2009).

45. Schindelin, J., Rueden, C. T., Hiner, M. C. & Eliceiri, K. W. The ImageJ ecosystem: An open platform for biomedical image analysis. Mol. Reprod. Dev. 82, 518–529 (2015).

46. Schneider, C. A., Rasband, W. S. & Eliceiri, K. W. NIH Image to ImageJ: 25 years of image analysis. Nat. Methods 9, 671–675 (2012).

